# STAMP: Single-Cell Transcriptomics Analysis and Multimodal Profiling through Imaging

**DOI:** 10.1101/2024.10.03.616013

**Authors:** Emanuele Pitino, Anna Pascual-Reguant, Felipe Segato-Dezem, Kellie Wise, Irepan Salvador-Martinez, Helena Lucia Crowell, Elise Courtois, William F. Flynn, Santhosh Sivajothi, Emily Soja, Sara Ruiz, Ginevra Caratù, Adrienne E. Sullivan, German Atzin Mora Roldan, B. Kate Dredge, Maycon Marção, Yutian Liu, Hannah Chasteen, Monika Mohenska, José Polo, Juan C. Nieto, Jasmine Plummer, Holger Heyn, Luciano Martelotto

## Abstract

We introduce Single-Cell Transcriptomics Analysis and Multimodal Profiling (STAMP), a scalable profiling approach of individual cells. Leveraging transcriptomics and proteomics imaging platforms, STAMP eliminates sequencing costs, to enable single-cell genomics from hundreds to millions of cells at an unprecedented low cost. Stamping cells in suspension onto imaging slides, STAMP supports single-modal (RNA or protein) and multimodal (RNA and protein) profiling and flexible, ultra-high-throughput formats. STAMP allows the analysis of a single or multiple samples within the same experiment, enhancing experimental flexibility, throughput and scale. We tested STAMP with diverse sample types, including peripheral blood mononuclear cells (PBMCs), dissociated cancer cells and differentiated embryonic stem cell cultures, as well as whole cells and nuclei. Combining RNA and protein profiling, we applied immuno-phenotyping of millions of blood cells simultaneously. We also used STAMP to identify ultra-rare cell populations, simulating clinical applications to identify circulating tumor cells (CTCs). Performing *in vitro* differentiation studies, we further showed its potential for large-scale perturbation studies. Together, STAMP establishes a new standard for cost-effective, scalable single-cell analysis. Without the need for sequencing, STAMP makes high-resolution profiling more affordable and accessible. Designed to meet the needs of research labs, diagnostic cores and pharmaceutical companies, STAMP holds the promise to transform our capacity to map human biology, diagnose diseases and drug discovery.

## Main text

Over the last decade, single-cell sequencing, particularly single-cell RNA sequencing (scRNA-seq), has revolutionized our understanding of complex tissues and organs by providing high-resolution maps of their individual cells. The profiling of single cells has offered invaluable insights into cellular composition and states during steady conditions and dynamic processes, such as differentiation or perturbations in diseases^1^. Innovations in microfluidics and combinatorial indexing have enabled the profiling of increasingly large cell numbers, generating atlases of entire organisms, including the fly, mouse and human^2–4^. Despite these advancements, current single-cell transcriptomic methods face significant limitations. The reliance on sequencing and the need to index cells individually in wells or droplets results in high costs that scale linearly with cell numbers. Additional limitations relate to the random sampling of cellular transcripts, resulting in a bias and overrepresentation of highly abundant transcripts at the cost of lowly expressed, often cell lineage defining genes, such as transcription factors.

Single-cell sequencing methods also suffer from inherent inefficiencies, with droplet-based microfluidics being affected by cell damage, inefficient encapsulation, and droplet instability, leading to significant sample loss. On the other hand, plate-based combinatorial indexing methods show limited cell capture, cross-contamination and inefficient indexing. Factors such as cell size, fragility and RNA content can further affect the capture efficiency, resulting in underrepresented or entirely missed cell populations^5^. As a consequence, current single-cell datasets may not accurately reflect the cellular composition and sample complexity.

Conventional scRNA-seq workflows, whether used alone or in combination with multimodal^6^ or multiplexing^7^ strategies have relatively low throughput, typically handling thousands of cells per experiment. Current methods face difficulties with both ultra-low (hundreds of cells) and ultra-high (millions of cells) experimental scales, the latter related to high costs^8^ and high input material^5,9^. Scaling single-cell profiling is crucial though for studying rare cell populations and to capture the full complexity of cell plasticity in health and disease. In addition, sequencing-based methods destroy cells during molecule capture and library preparation, limiting the ability to combine molecular profiles with cell morphology (e.g., shape and size) or functional attributes (e.g. metabolic activity)^10^.

To address these limitations, we hypothesized that imaging-based transcriptomics and proteomics readouts could significantly scale-up cell numbers at substantially reduced costs, while preserving the advantages of molecule detection at a single-cell level. Advances in spatial transcriptomic imaging assays now offer large gene panel designs, operating at transcriptome-wide scale, closely mirroring the capabilities of scRNA-seq methods. Traditionally, spatial transcriptomics and proteomics have been applied in the context of tissue profiling to map the composition and architecture of complex samples, such as organs or tumors^11–13^. In this study, we transform two imaging platforms, the CosMx Spatial Molecular Imager (SMI, Nanostring Technologies/Bruker) and the Xenium Analyzer (10x Genomics), into scalable and flexible single-cell profiling tools. We termed this approach Single-Cell Transcriptomics Analysis and Multimodal Profiling (STAMP) through imaging, an adaptation of single-molecule imaging that enables the analysis of hundreds to millions of individual cells in suspension at unprecedented flexibility and scale. STAMP designs include single-modal (RNA or protein) or multimodal (RNA and protein) profiling, in single or multi-sample configurations, demonstrated across a range of sample types and experimental scenarios.

We evaluated STAMP’s scalability by profiling a broad spectrum of cell numbers, from hundreds to millions of cells, showcasing its ability to handle diverse sample sizes. We then tested the ability to profile both whole cells and nuclei, demonstrating its versatility in handling samples with different RNA contents and offering the potential to profile archived tissues^14,15^. We next applied both single-modal and multimodal profiling in the context of immuno-phenoptyping of millions of blood cells. We also used STAMP to simulate the identification of ultra-rare cell populations, such as CTCs, which are challenging to detect using conventional scRNA-seq methods. Finally, we applied STAMP to *in vitro* models, specifically, the differentiation of human embryonic stem cells (hESCs) with bone morphogenetic protein 4 (BMP4) as a model of gastrulation, illustrating its utility for capturing cell lineage architecture in complex developmental systems.

## Results

### Sequencing-free single-cell genomics through imaging

To address current limitations in single-cell genomics, we developed STAMP, a flexible and scalable approach for cost-efficient, massive-parallel, single-cell profiling. STAMP enables the profiling of single cells in suspension on slides integrated with state-of-the-art transcriptomic and protein imaging platforms, such as the CosMx SMI or the Xenium Analyzer. STAMP begins with the fixation and permeabilization of cells in suspension, followed by anchoring (“stamping”) cells onto instrument-compatible glass slides to form monolayers. Flexibility is achieved through the adaptable format and loading of stamping areas, allowing versatile multi-sample profiling. After stamping, the cells undergo gene set probe or oligo-conjugated antibodies hybridization and cyclic decoding through imaging, which can accommodate any gene or protein panel size and design **(Fig. 1a)**. This versatility allows STAMP to be used for data-driven, hypothesis-free strategies to explore the complexity of hundreds to millions of cells in a single experiment.

**Figure 1.**
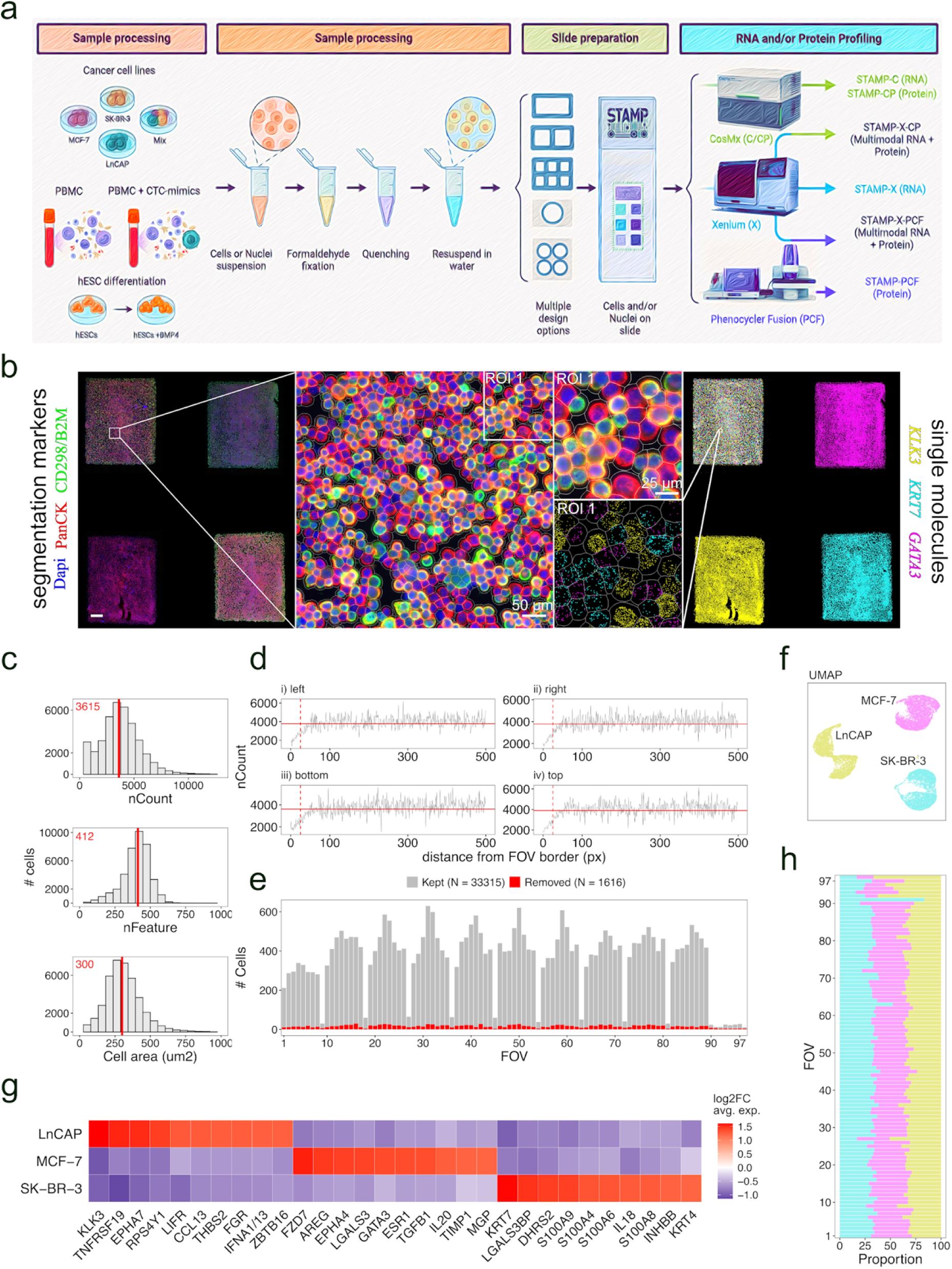
STAMP Experimental Design and Data Analysis Workflow. (a) Schematic of the STAMP workflow. STAMP-XPCF and STAMP-PCF data not included in this preprint. (b) Immunofluorescence (IF) image of STAMP-C showing DAPI staining (blue), CD298/B2M (red), and pan-cytokeratin (PanCK, green) across the four sub-STAMPs, with an enlarged view of an exemplary field of view (FOV). Marker genes specific to each cell line are highlighted for the entire STAMP and a selected region of interest (ROI): KLK3 for LnCAP in yellow, GATA3 for MCF-7 in magenta, and KRT7 for SK-BR-3 in cyan (each dot represents a transcript). Scale bar = 750 µm unless otherwise noted. (c) Quality metrics of the mixed sub-STAMP (containing equal representation of the three cell lines: MCF-7, LnCAP, and SK-BR-3) showing distributions and median values of counts, features, and cell area before filtering. (d) Line plot displaying the total number of counts relative to the distance from each FOV border. The horizontal red lines indicate the moving median, while the vertical dashed red line shows the applied threshold. (e) Bar plot illustrating the number of cells removed from each FOV due to the border effect. (f) UMAP of the mixed sub-STAMP (as shown in c), colored by InSituType unsupervised clustering. (g) Heat map showing the top 10 marker genes for each cluster identified in (f), representing each cell line, and normalised by feature. (h) Composition of each of the 99 imaged FOVs in the mixed sub-STAMP (as shown in e), indicating the distribution of cell lines.

To evaluate STAMP’s suitability and specificity for imaging-based transcriptomics, we profiled three cancer cell lines (LnCAP, MCF-7, and SK-BR-3) using the CosMx SMI platform (STAMP-C) with the 1000-plex Human Universal Cell Characterization RNA Panel. The multi-sample slide array contained four sub-STAMPs, each containing ∼35,000 cells: three with individual cancer cell lines and one with a 1:1:1 pooled mixture **(Supplementary Fig. 1a)**. For imaging, we selected contiguous fields of view (FOVs) to comprehensively scan each sub-STAMP. We also obtained cell staining information for DAPI, pancytokeratin (PCK) and pan-membrane markers (B2M/CD298), used for cell segmentation (see Methods; **Fig. 1b**). We initially confirmed a uniform distribution of transcripts, genes and cell areas across each sub-STAMP pointing to homogeneous cell stamping (**Supplementary Fig.1c-d**). We then tested the pooled mixture to distinguish different cell types in a spatially mixed environment. On average, we obtained 11,103 cells per cell line, with a median of 3,637 transcripts and 413 genes per cell **(Fig. 1c)**. Stringent quality control steps included removing segmentation artifacts, filtering out low transcript and gene counts (<2.5 median absolute deviations, MAD), and excluding high counts and large cell areas (>2.5 MAD). Cells near the FOV borders (<30 pixels) were also excluded due to reduced counts and features **(Fig. 1d)**. Overall, 4.63% of cells were removed from the analysis **(Fig. 1e)**.

Using the InSituType (IST)^16^ algorithm for unsupervised clustering, we identified three distinct clusters with unique transcriptional profiles **(Fig. 1f,g)**. The pooled mixture consistently contained an average of 33.33% (range: 32.6 - 33.4%) of each cell line per FOV **(Fig. 1h)**. To validate the clustering results of the pooled mixture, IST was applied to the sub-STAMPs of the individual cell lines, which also separated into three distinct clusters with unique transcriptional profiles matching those found in the pooled mixture **(Supplementary** Fig. 1e,f). The gene signatures of both pooled and individual cell lines were correlated with data from a scRNA-seq reference dataset generated using the Single Cell Gene Expression Flex assay (10x Genomics) on the same suspension of fixed cancer cells **(Supplementary** Fig. 1g).

### Sensitive capture of low-input cell numbers

To explore the scalability to ultra-low cell densities, we stamped cancer cell line mixtures (1:1:1 ratio of MCF-7, SK-BR-3, and LnCAP) into four sub-STAMPs with varying cell counts of approximately 100, 250, 500, and 1,000 cells (**Fig. 2a**). This setup allowed us to assess the performance at low cell numbers, which are challenging to capture with current droplet microfluidics or combinatorial indexing scRNA-seq methods. Additionally, we profiled ∼20,000 cells and ∼20,000 nuclei of the cell line mixture separately in two sub-STAMPs, enabling a direct side-by-side comparison of high-RNA (whole cell) and low-RNA (nuclear) content samples within a multiplexed experimental setup. Again, contiguous FOVs were selected, ensuring comprehensive coverage of the low-density samples.

**Figure 2.**
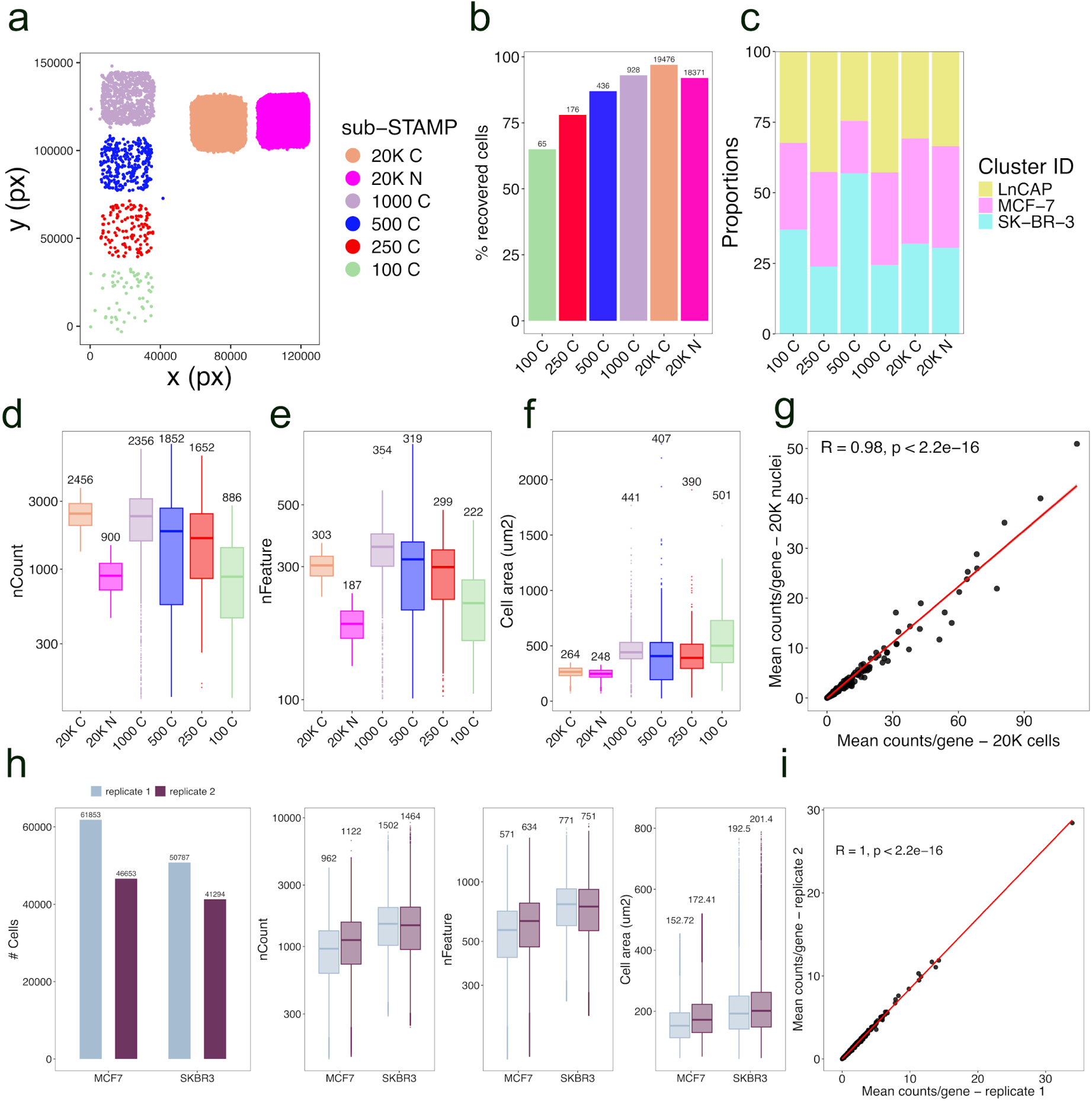
STAMP is a robust, scalable and flexible platform for imaging-based single cell transcriptomics of cells in suspension. (a) Spatial plot showing the experimental design of STAMP-C with 6 sub-STAMPs to showcase low numbers of tumor cells seeded (sub-STAMPs 100 C, 250 C, 500 C and 1,000 C and sub-STAMPs 20K C and 20K N for cells and nuclei comparison). Each dot is a cell, as detected by imaging. b) Percentage of cells retrieved from input cells across different sub-STAMPS. Text labels indicate absolute cell counts. (c) Proportions of tumor cell lines across sub-stamps. Box plot displays the number of counts (d), features (e), and cell area (f) for each sub-STAMP with respective medians. (g) Pearson correlation of raw counts averaged by cell number between cells and nuclei. (h) Number of detected (segmented) cells, number of detected transcripts and genes, and cell area for each cell line in each STAMP-x (replicate 1 and 2). (i) Pearson correlation of raw counts averaged by cell number between MCF-7 and SK-BR-3 replicates. (g, i) Each dot is a gene. Red line shows the fitted linear regression

The mean recovery rate of high-quality cells was 85.33% (range: 65 - 97 %) from the input material (**Fig. 2b**), with an average mix of 32.5% for each cell line per sub-STAMP (range: 18.6 - 56.9%, **Fig. 2c**). The median gene counts per cell were consistent across sub-STAMPs representing higher loadings, with reduced genes detected in the lowest sub-STAMP (100 cells; **Fig. 2d**). The number of detected features per cell followed similar trends, indicating suboptimal performance when profiling extremely low cell densities (**Fig. 2e**). Cell areas remained stable in less densely seeded sub-STAMPs, but halved in those with higher confluence, likely due to reduced cell expansion during segmentation (**Fig. 2f**). Expectedly, nuclei had fewer counts and features than intact cells, yet, gene expression levels were highly correlated across cells and nuclei **(Fig. 2d,e,g)**.

We next tested the reproducibility of our approach by splitting MCF-7 and SK-BR-3 cell suspensions into two aliquots, imaged in two slides as technical replicates (scanned in the same Xenium run using the Xenium Prime 5k Human Pan Tissue and Pathways panel, STAMP-X). While recovered cell numbers slightly varied, both replicates showed a highly consistent number of transcripts and genes detected, as well as equal cell areas (**Fig. 2h**). Accordingly, gene expression levels highly correlated across replicates (**Fig. 2i**).

### Immuno-phenotyping of millions of circulating blood cells

We then expanded the protocol to accommodate millions of cells per experiment. We generated a high-density STAMP containing ∼1.7 million PBMCs, on which we applied the Xenium Immuno-Oncology panel (380 target genes). Imaged cells showed a median of 83 transcripts per cell (range: 27 - 259) and 49 genes per cell (range: 24 - 103). After quality control to remove segmentation errors and cells with extreme counts and features (**Supplementary** Fig. 2), we retained 88,53% of high-quality cells. We next performed scRNA-seq analysis strategies, including dimensionality reduction and principal component analysis (PCA, see **Methods**). PCs 1-3 were sufficient to accurately resolve the three main immune lineages (i.e. Myeloid, T and B cells) with cell lineage marker genes driving the separation (**Fig. 3a,b**). Further clustering of each immune cell lineage led to the identification of all major PBMC cell types and states at expected frequencies and cell areas, represented by cell state defining markers (**Fig. 3c,d,e**). In detail, we identified 13 different immune clusters, representing the main populations of immune cells in PBMCs. To further test the limits to which STAMP can be used for fine-grained immuno-phenotyping, we stamped another 750,000 cells, profiled with a larger gene probe panel (Xenium Prime 5k Human Pan Tissue and Pathways panel). Strikingly, zooming into the main immune cell lineages allowed us to generate a high-resolution map of 31 immune cell states in circulation, the basis for large-scale atlasing projects across different dimension, such as time (e.g. age) or genetics (ethnic background), but also suitable for multiplexed perturbation studies (**Fig. 3f,g**). In detail, we identified eight CD4+ T cell subsets, spanning from naive to effector and memory populations and capturing the Th1, Th2 and Th17 functional cell types that define the three arms of adaptive cell-mediated immune responses. Similarly, we found eight subclusters within the CD8+ T cell pool: from naive to central/effector memory, as well as different types of effector populations, including interferon-responding CD8 T cells. Within the circulating NK cell pool, we distinguish CD56^dim^CD16^bright^ from CD56^bright^CD16^dim^ NK cells, two subsets with distinct antibody-dependent cellular cytotoxicity and divergent migratory properties, together with a small proportion of proliferating *EOMES*- and *CD34*-expressing cells, which might represent NK progenitors. Ig-related genes and other markers relevant for B cell phenotyping were missing in the panel, which limited a deeper annotation of B cell differentiation states. However, the myeloid compartment showed the well-defined monocyte populations found in blood (classical, intermediate and non-classical monocytes) and allowed us to define dendritic cells with granularity^17^.

**Figure 3.**
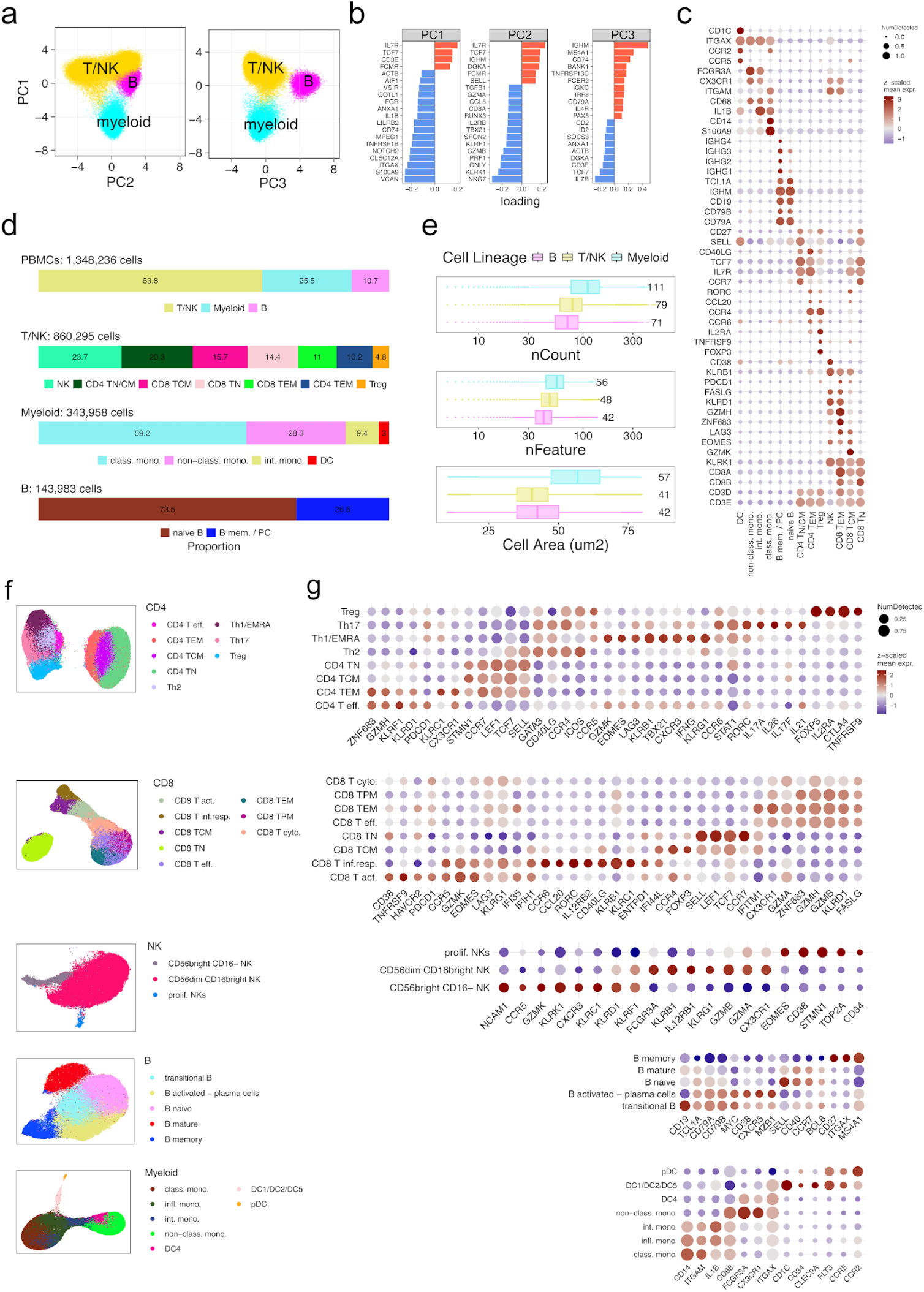
Profiling ultra-high cell numbers with STAMP-X. (a) Principal component analysis (PCA) color-coded by cluster identity, based on Leuven clustering of the PBMC dataset analyzed with the Immuno-oncology panel. (b) Loadings of the 3 first PCs. (c) Dotplot showing cell type defining markers for each cluster normalised by feature after sub-clustering of main immune lineages as in (a). See methods for further details. (d) From top to bottom, total cell numbers and proportions of the whole PBMC dataset and of each cell lineage as defined in (a). (e) Box plot display of the number of counts, features and cell area split by cell lineage. Text labels indicate median values. (f) UMAPs of the three main immune lineages from the PBMC dataset analyzed with the Xenium Prime 5k Human Pan Tissue and Pathways panel, colored by cell type. (g) Dot plot showing cell type-defining marker genes normalized by feature for each cluster identified in (f). DC, dendritic cells; pDC, plasmacytoid dendritic cells; non-class. mono; non-classical monocytes; int.mono; intermediate monocytes; infl. mono.; inflammatory monocytes; class.mono., classical monocytes; B mem./PC, memory B cells / plasma cells; TN, naive T cells; TCM, central memory T cells; TEM, effector memory T cells; TPM, peripheral memory T cells; T eff., effector T cells; T act., activated T cells; T cyto.; cytotoxic T cell; T inf.resp.; interferon responder T cells.

### Combined RNA and protein multimodal profiling

Having established an imaging-based approach for scalable and cost-efficient single-cell transcriptomics, we next tested the possibility to combine RNA and protein readouts within the STAMP framework. To this end, we used the slides from the previously described transcriptomic experiment performed on the Xenium and subsequently applied protein profiling on the CosMx using the 64 protein panel (STAMP-X-CP). In parallel, we performed a standard CosMx protein experiment to compare how the prior STAMP-X application affects the outputs (STAMP-CP). Both conditions resulted in high-quality protein profiles, with high protein counts per cell (**Fig. 4a**). We noticed, though, that the CosMx protein profiling alone showed higher cell areas, DAPI mean fluorescence intensity and protein counts, pointing to increased quality when running protein analysis alone (**Fig. 4a,b**). Nevertheless, protein levels strongly correlated between the experiments, indicating robust protein detection in the multimodal STAMP conditions (R=0.93; **Fig. 4c**). Most importantly, the protein data could be integrated between both experiments and used for protein-based immuno-phenotyping (**Fig. 4d,e**). Given increasing protein panel sizes, such multimodal STAMP readout will allow for data-driven analysis integrated for both RNA and protein information, beyond the here highlighted use of single-modality analysis.

**Figure 4.**
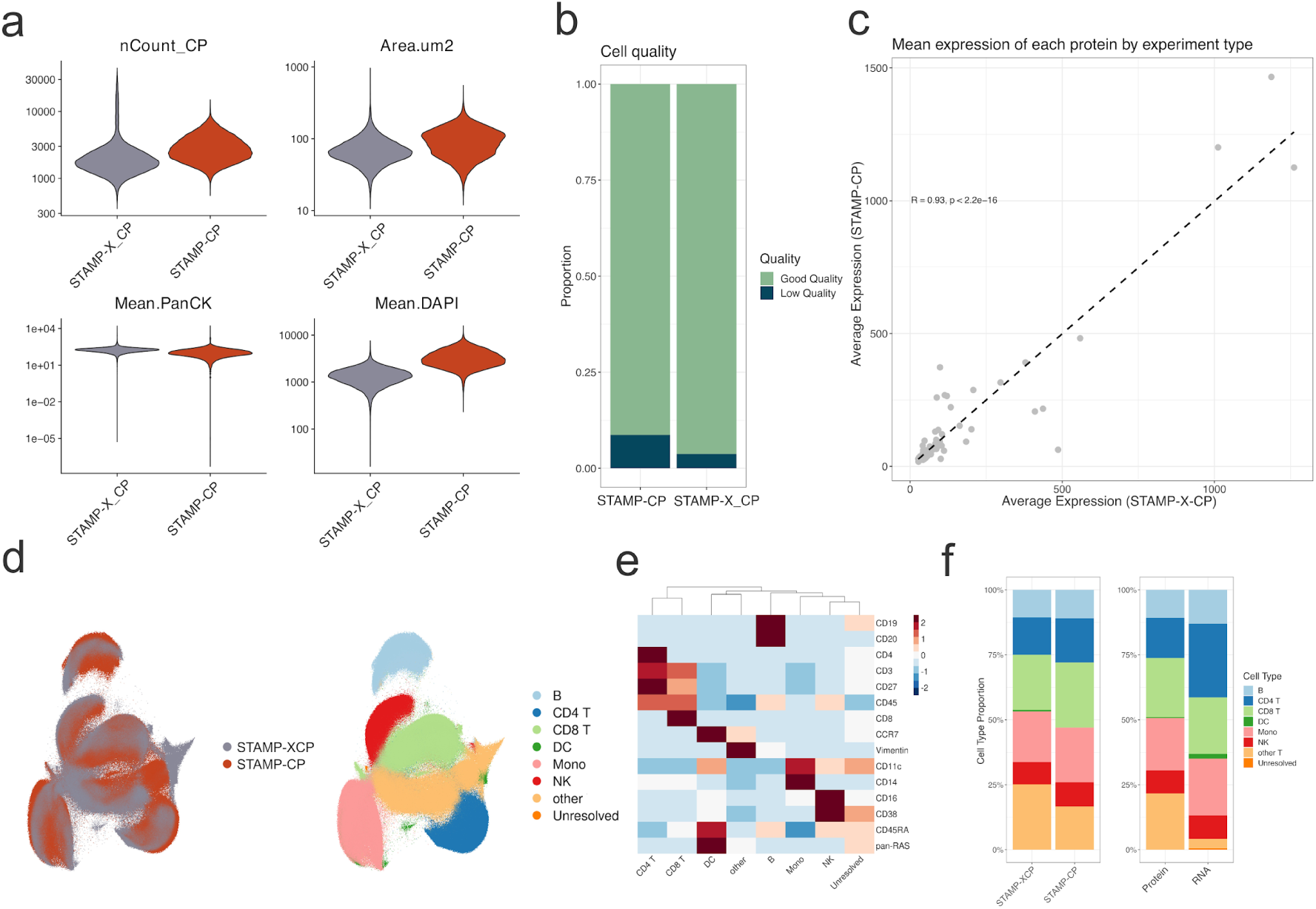
STAMP Protein and multimodal experiment QC. (a) Violin plots showing four key metrics used for quality control across differently run experiments. STAMP-X-CP was run using CosMx Protein after Xenium RNA, while STAMP-CP was run using CosMx Protein alone. (b) Bar plot depicting the number of cells yielded in each experiment, with bars coloured according to quality. (c) Scatter plot where each dot (n = 62) represents a protein in the CosMx panel; the X-axis shows the average expression of these proteins in the STAMP-X-CP experiment, and the Y-axis shows the average expression in the STAMP-CP experiment. The line represents the best fit through the data points using a linear model. R and p-value statistics are provided for Pearson correlation analysis. (d) UMAP plot illustrating the dimensionality reduction of both experiments on the same embedding, showing unsupervised clustering and cell type annotations. (e) Average expression heatmap of protein markers used for coarse cell type annotation, normalized expression values were aggregated for all cells and scaled across features.(f) Stacked bar plots showing: (i) the proportion of each cluster in STAMP-X-CP and STAMP-CP, (ii) the proportion of each cell type in STAMP-X-CP and STAMP-CP, and (iii) the proportion of cell types per modality of the same cells run in STAMP-CP and STAMP-X.

### Sensitive detection of rare cell types

Given the scalable design of STAMP, we were next interested in simulating clinically relevant scenarios, in particular, the identification of CTCs. Tumor cells circulating in the blood provide a valuable measure to quantify tumor burden, but also to identify tumor heterogeneity and actionable alterations. To simulate CTC detection from blood samples and to determine sensitivity limits of the protocol in this context, we spiked-in MCF-7 cancer cells at 1:100,000 and 1:50,000 dilutions into PBMC samples, stamped onto the same slide (total 1.1 million cells). We then imaged the cell mixtures with the CosMx Human Universal Cell Characterization RNA panel. The CosMx cell segmentation markers already allowed for the manual detection of the CTC-mimics, which were randomly distributed across each sub-STAMP (PCK staining, **Fig. 5a**). To automate CTC-mimic detection, we generated gene expression signatures using scRNA-seq performed on fixed cells and scored each imaged single cell based on their MCF-7 or PBMC profiles **(Fig. 5b)**. We additionally combined information of the PCK mean fluorescence intensity, the number of transcript counts and the cell areas, to jointly identify CTC-mimic with high accuracy **(Fig. 5c,d)**. Thereby, in sub-STAMPs with 10 and 20 target spike-in cells, we confidently identified 7 and 28 CTC-mimics, respectively. Here, CTC-mimic accounted for 0,001% of the overall sample size, demonstrating the capacity to detect extremely rare cellular types with clinical application potential. Of note, the immuno-fluorescence images allowed us to manually validate the CTC-mimic identities based on their coordinates registered with the STAMP experiments (**Fig. 5e)**. The detection of these rare events is independent of the platform, panel, or segmentation method, as demonstrated by the identification of CTC-mimics using Xenium with the Xenium Prime 5k Human Pan Tissue and Pathways panel (**Supplementary** Fig. 3).

**Figure 5.**
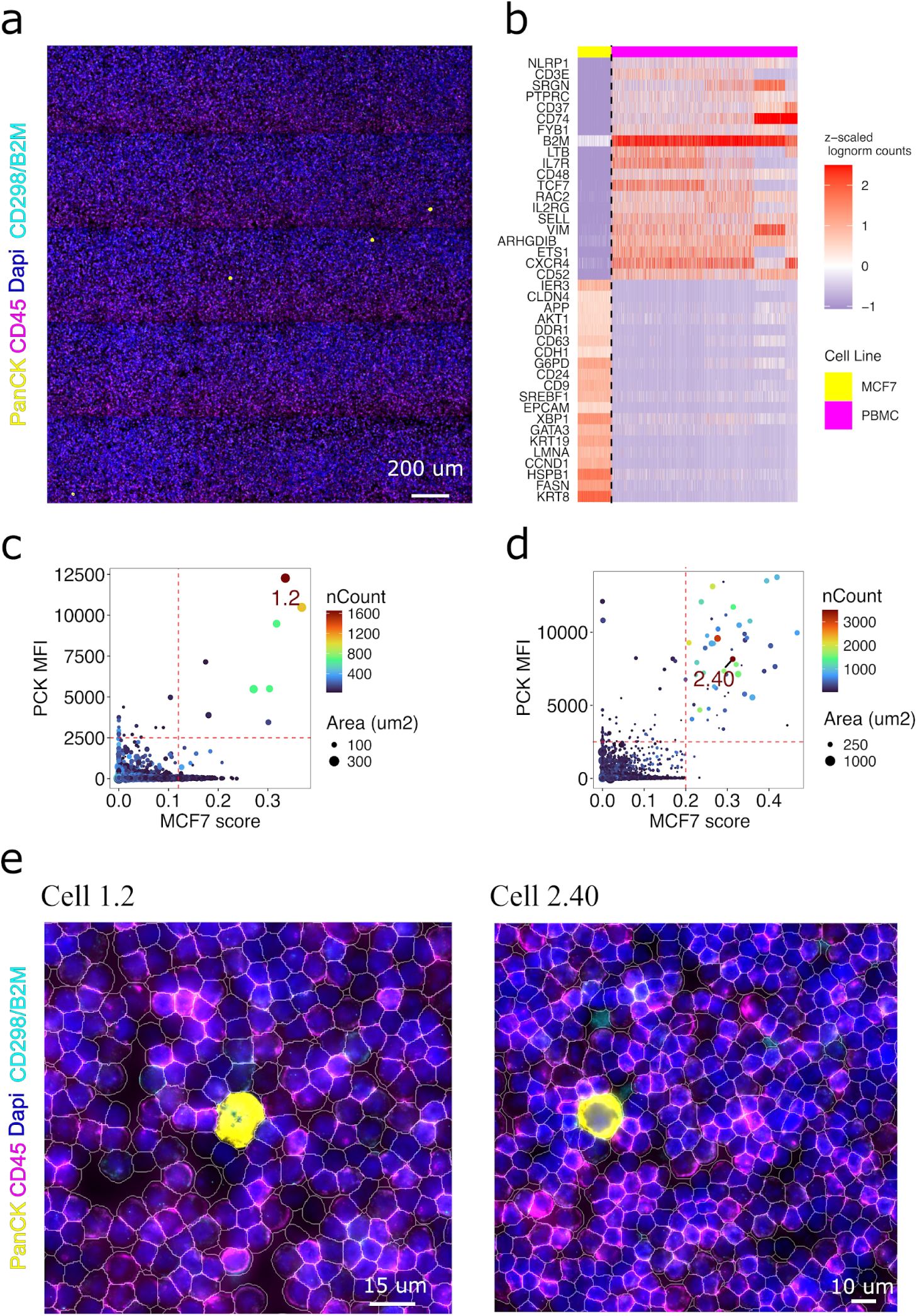
Detection of circulating tumor cells mimics (CTC-mimics) with STAMP. (a) Immunofluorescence (IF) image of an exemplary field of view of the STAMP with 10 CTCs, showing DAPI in blue, pancytokeratin (PanCK) in yellow, the pan membrane marker CD298/B2M in cyan and CD45 in magenta. (b) Heat map of the MCF-7 and PBMC gene signatures extracted from the differential expression analysis in the Flex dataset subsetted for the CosMx 1K panel. (c) Scatter plot shows the MCF-7 signature score, as extracted in (b) and PanCK mean fluorescence intensity (MFI) in the 10 CTC-mimics spike-in substamp. Each dot is a cell, with size proportional to cell area and color-coded by the number of counts. (d) same as B but in the 20 CTC-mimics spike-in substamp. (e) IF images of two regions of interest showing DAPI in blue, pancytokeratin (PanCK) in yellow, the pan membrane marker CD298/B2M in cyan and CD45 in magenta and the cell segmentation outlines in white, where DAPI+ PanCK+ CD45- CTC-mimcs #1.2 identified in (c) and #2.40 identified in (d) are depicted.

### Profiling continuous cell states during stem cell differentiation

We next tested the resolution to which we can identify cell states emerging from differentiating hESCs in response to BMP4 treatment, leveraging the multiplexing capabilities of STAMP. Adding BMP4 to cultured hESCs initiates signaling cascades similar to those observed in the epiblast during gastrulation, resulting in a variety of embryonic and extra-embryonic cell types. We combined eight BMP4 treatment timepoints (0 to 120 hours) in a single STAMP experiment using the CosMx Human Universal Cell Characterization RNA panel (**Fig. 6a,b**). We then performed pseudotime trajectory analysis using Palantir to model the differentiation path of hESCs in response to the BMP4 treatment (**Figure 6c-f**). The hESCs treated with BMP4 progressed from a pluripotent state to an intermediate state after 12/24h of BMP4 treatment, marked by a downregulation of *SOX2* and upregulation of *GATA3* expression. By 48h a Mesendoderm-like state appears, expressing *EOMES* and *KDR*, together with an Amnion precursor-like state, strongly expressing *GATA3*. By 72h the Mesendoderm-like state bifurcated into Early Mesoderm-like and Endoderm-like cell states, marked by the expression of *SNAI2* and *PDGFRA* or *CXCR4* and *APOA1*, respectively. After 72h, Amnion precursor cells differentiate into more mature Amnion-like cells expressing *TGFBI*, while the Mesoderm branch differentiates further into a Late Mesoderm-like cell state showing upregulation of *DUSP6* and *FOXF1*. Using Palantir, we also computed the gene expression dynamics of key marker genes along the Amnion, Mesoderm and Endoderm branches, showing the progressive downregulation of pluripotent genes and upregulation lineage defining markers (**Figure 6g**).

**Figure 6.**
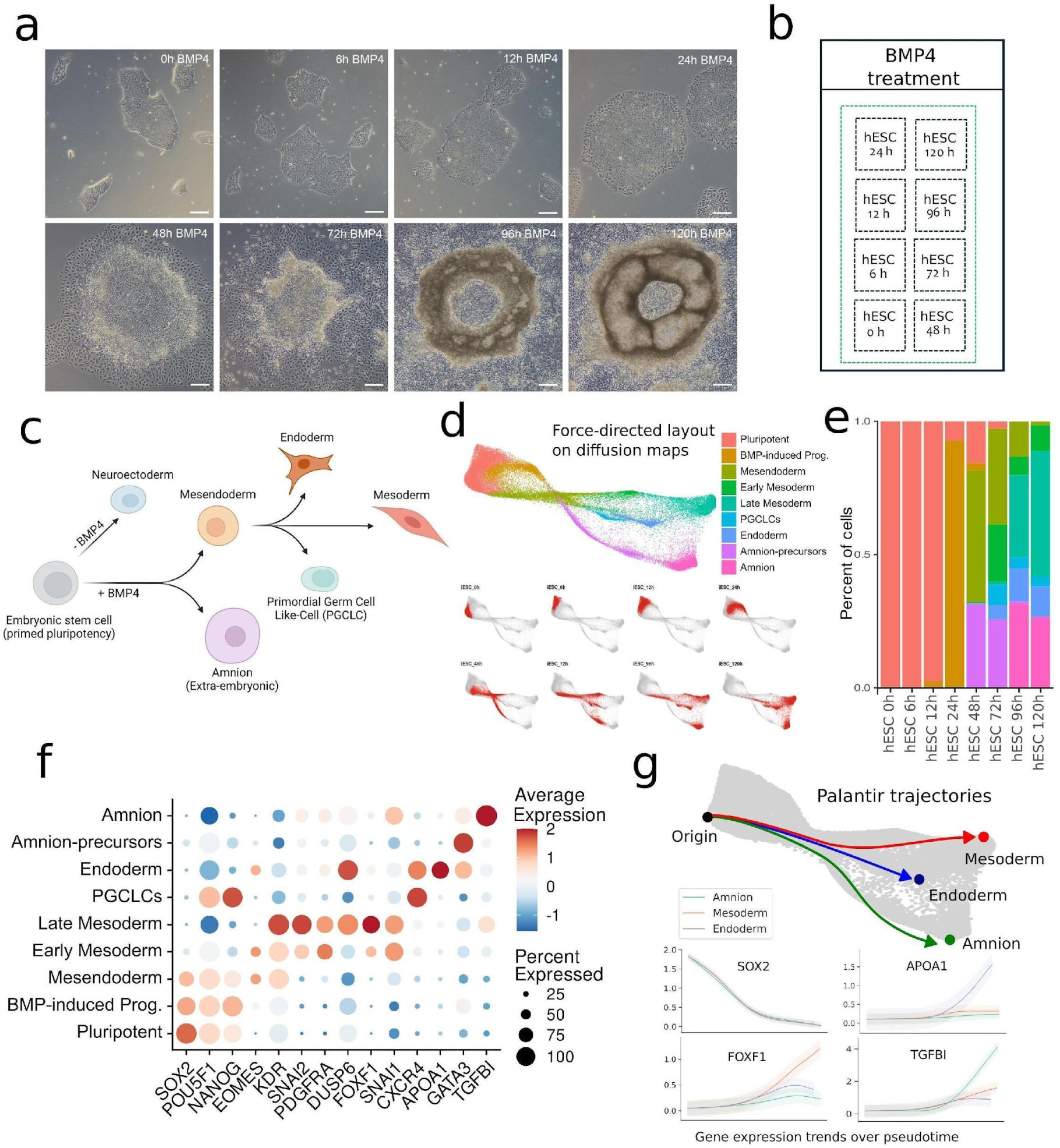
Profiling hESCs differentiation after BMP4 induction. a) H1 (WA01) human embryonic stem cells treated with BMP4 (50 ng/ml) for 0, 6, 12, 24, 48, 72, 96, 120 hours. Images captured on a Zeiss Primovert microscope. Scale bars show 200 µm. b) Layout of the STAMP slide with the 8 timepoints after BMP4 treatment. c) Schematic showing the expected cell states after BMP4 treatment of hESCs. d) Cells from the eight timepoints after BMP4 treatment projected on a force-directed layout on diffusion maps computed with the Palantir algorithm. On top cells are colored based on the cell state annotation and below cells from each timepoint are shown separately. e) Proportion of annotated cell states in each of the timepoints. Color code is the same as panel d. f) Dotplot showing the main marker genes used for the cell state annotation. g) The differentiation trajectories for Amnion, Endoderm and Mesoderm were inferred with Palantir (top), as well as expression trends for selected marker genes along the trajectories’ pseudotime (bottom).

## Discussion

We evaluated STAMP with the main objectives to verify: 1) specificity, 2) reproducibility, 3) scalability, 4) sensitivity, 5) resolution and 6) flexibility. Cancer cell lines were used to confirm specificity and to identify potential artifacts, ensuring that STAMP accurately targeted specific gene sets within individual cells without cross-hybridization or background noise. For reproducibility and scalability, we tested the analysis of millions of PBMCs across different platforms, demonstrating STAMP’s consistent performance. Flexibility was assessed using diluted PBMC samples and multiplexing experiments, proving the adaptability in handling varying cell numbers (from hundreds to millions of cells) and multiple samples simultaneously (e.g., *in vitro* differentiation models). To showcase the high cellular resolution, we immuno-phenotyped blood cells to high granularity and analyzed cellular states during differentiation. The sensitivity was tested using CTC-mimic experiments, ensuring STAMP could detect and characterize very lowly abundant cell populations.

STAMP provides a unified solution that integrates high-throughput analysis, cost efficiency, and multi-sample profiling, expanding the possibilities of single-cell analysis for diverse research and clinical applications, and for advancing our understanding of complex biological systems and diseases. STAMP transforms imaging-based genomics platforms into high-throughput profiling tools for cells in suspension. The versatile platform supports single-modal (RNA or protein) and multimodal (RNA and protein) profiling, bridging the gap between mRNA and protein expression levels. STAMP also offers unprecedented flexibility in experimental design, including the simultaneous analysis of multiple samples and conditions. By eliminating sequencing costs, STAMP establishes a new standard in single-cell research, making high-resolution, cost-effective profiling accessible to a broader community, including core labs, academic institutions, and pharmaceutical companies. STAMP’s ability to detect rare populations with high sensitivity and specificity, such as CTCs in minimal residual disease, makes it a powerful tool in clinical genomics and precision medicine.

Its ability for visual and molecular profiling without sequencing, sets a new benchmark for scalable, high-throughput single-cell analysis. Unlike sequencing-based methods, STAMP leverages advanced imaging techniques that allow each cell to be spatially indexed, capturing morphological information alongside molecular data. Imaging individual cells allows researchers to visually inspect cells, as shown here for CTC-mimics, particularly valuable in the clinics where histopathology validation is standard-of-care. Moreover, cross-referencing molecular data with visual characteristics, reduces false positives to improve reliability. Additionally, the non-destructive nature of STAMP allows for simultaneous analysis of RNA, protein, and histological features of the same cells, enabling comprehensive multimodal profiling to support multiscale phenotype readouts. The ability of STAMP to handle a wide range of cell numbers at low cost allows researchers to tailor experiments, focusing on small, rare cell populations, conducting large-scale cell atlases or CRISPR and drug screens.

STAMP addresses current limitations in single-cell genomics by offering a solution that combines scalability, cost-effectiveness and the flexibility, within a single approach. This allows researchers to conduct experiments with a broad range of sample sizes, from limited biopsy specimens to extensive cellular atlasing, without compromising data quality or increasing costs. This versatility makes STAMP an invaluable tool for a wide range of experimental needs, from basic research to clinical applications.

## Methods

### Maintenance of Cancer Cell Lines

MCF-7 cell line was cultured in Dulbecco’s Modified Eagle Medium (ThermoFisher Scientific, 11885084) containing 10% Fetal Bovine Serum (FBS) and 2 mM L-Glutamine. SK-BR-3 and LnCAP cell lines were grown in McCoy (modified) 5A medium (Thermo Fisher Scientific, 16600082) and RPMI-1640 medium, respectively, both supplemented with 10% FBS (ThermoFisher Scientific). All cultures were maintained at 37 °C, 5% CO₂. Cells were trypsinized using TrypLE™ Express Enzyme (ThermoFisher Scientific, 12604013) and cryopreserved in freezing media composed of 90% FBS and 10% DMSO (Merck, 41639). Cells were frozen in a CoolCell Container (Corning) at -80 °C overnight at a density of 1 × 10⁶ cells/mL and transferred to -150 °C storage the following day. Cell counting and viability measurements before and after storage were performed using the Luna-FX7 Automated Cell Counter (Logos Biosystems, AO/P Viability Kit, F23011). Cell viability after thawing was >85%. On the day of fixation, cells were thawed in a 37 °C water bath, washed twice with PBS (without Ca²⁺/Mg²⁺) + 0.05% BSA (Miltenyi Biotec, 130-091-376).

### Differentiation of Human Embryonic Stem Cell (hESC)

Human embryonic stem cells (hESCs, WA01, WiCell) were cultured in vessels coated with growth factor-reduced Matrigel (Corning, 354230) and maintained in mTeSR+ growth media (STEMCELL Technologies, 85850). Cultures were passaged by removing the medium, washing with DPBS (Sigma-Aldrich, D8537), and treating with Gentle Cell Dissociation Reagent (STEMCELL Technologies, 100-0485,) at sufficient volume to cover cells for 3 min at 37 °C. The dissociation reagent was aspirated, growth media was added, and cells were lifted by scraping with a cell lifter to produce small cell clumps. One-sixth volume of lifted cells was added to a new culture vessel and topped up with mTeSR+ growth media. All cultures were grown at 37 °C, 5% CO₂. For differentiation, hESCs were seeded as colonies into 6-well trays by passaging as described. The following day, hESCs were treated with 50 ng/mL human recombinant BMP4 (STEMCELL Technologies, 78211) in mTeSR+ media for 0, 6, 12, 24, 48, 72, 96, or 120 h, with daily media replacement. To harvest, media was removed, cells were washed with DPBS, and Gentle Cell Dissociation Reagent was added to cover the cells. Cultures were incubated at 37 °C until dissociated, then growth media was added. Dissociated cells were pelleted by centrifugation (5 min, 200 × g), and the media was aspirated. Cell pellets were resuspended in 1 mL mFreSR (STEMCELL Technologies, 05855) and transferred to a cryotube, then frozen in a CoolCell Container (Corning) at -80 °C overnight. Cells were transferred to -150 °C storage the next day. Cell counting and viability measurements before and after storage were carried out using the Luna-FX7 Automated Cell Counter. Viability of cells after thawing was >85%.

### Slide Preparation and ‘STAMPing’ Procedure

Superfrost Plus Micro Slides (VWR, 48311-703) for STAMP-C/CP/PCF and Xenium slides (10X Genomics, PN-3000941) for STAMP-X/X-CP/X-PCF/PCF, were placed on Xenium slide holders were coated with 1 mL of Poly-D-Lysine for 1 h and overnight at 37 °C, respectively, in a thermocycler using the Xenium Thermocycler Adapter plate positioned atop the 96-well block of a C1000 Touch Thermal Cycler (BioRad) with the lid closed and set at 37 °C. After coating, the slides were washed with 1 mL of Nuclease Free Water (ThermoFisher Scientific, 10977023) 3 times and air dried. Custom single or multi-plex (multi-sample) arrays of various areas/volumes were created using either 10X Genomics’ gaskets (PN- 370017) from the Chromium Single Cell Reagent Kits and using a hole punch or punch pliers (Total Tools, 9070220SB), micro-Slide 8-well (ibidi, 80841) and 12-well (ibidi, 81201) cell culture chambers or a silicone gasket for ProPlate® microarray system (Grace Bio-Labs, 246875) by placing them on the coated slides within the scanning area of the CosMx, Xenium or Phenocycler. For the STAMPing procedure (i.e. attaching cells on slides for STAMP), up to 5 million single cells in suspension, with >80% viability, were first fixed with 4% Formaldehyde (Sigma Aldrich, 252549-500ML) + 1x Concentrated Cell Fixation and Permeabilization Buffer (10x Genomics, PN- 2000517), as per the Fixation of Cells & Nuclei for Chromium Fixed RNA Profiling protocol (CG000478, RevD). After 2 h incubation at RT, cells were pelleted at RT for 5 min at 850 rcf and resuspended in 1 mL of 1x Quenching Buffer (10x Genomics, PN-2000516), washed 2 times with nuclease-free water and finally resuspended in water + 0.01% Triton-X and counted in duplicates or triplicates using the Luna-FX7 Automated Cell Counter. To maximize the use of the area in each stamp (or substamp, when in multi-sample settings) for high and ultra-high cell profiling we used the average cell size data provided by the Luna-FX7 Automated Cell Counter and estimates generated using ChatGPT 4o to calculate the number of cells that would fully (or partially) cover a desired area of the given custom array. Before STAMPing, cell concentrations were adjusted so that the minimum volume of cell suspension added to the wells would evenly cover the bottom surface for the desired number of cells to profile and fully dry within 1 h of incubation at 42 °C. This volume was dependent on the well area and was approximated in advance for each well size. Cells were then loaded into the wells, and the slide was placed in a thermocycler using the Xenium Thermocycler Adapter plate, running the following program: 4 °C for 30 min, 25 °C for 5 min, 42 °C for up to 1 h (or until the volume dried), followed by 42 °C for 2 h, and then held at 22 °C. After STAMPing, custom wells were carefully removed, and the slides were either processed immediately or placed in a mailer with desiccant at RT for STAMP-X/X-CP/X-PCF or at 4 °C for STAMP-C/CP/PCF until further processing. The same procedure was carried for STAMP profiling using PBMCs from a healthy donor (STEMCELL Technologies, 200-0470), cancer cell lines (MCF-7, SK-BR-3 and LnCAP, either individually or mix 1:1:1 MCF-7:SK-BR-3:LnCAP), iPSC, hESC and CTC-mimics. For CTC-mimics, MCF-7 and SK-BR-3 cell lines were counted in triplicates and spiked into 1 million PBMCs either as a single line (MCF-7) or as a mix (1:1 MCF-7:SK-BR-3). Spike-in ratios varied from less than 10 to more than 10 but fewer than 50 cancer cells per approximately 1 million PBMCs.

### CosMx RNA Slide Preparation (STAMP-C)

STAMPed slides were placed on a Xenium Thermocycler Adapter plate atop the 96-well block of a thermocycler with the lid open and incubated at 60 °C for 2 h, then equilibrated to RT for 3–5 min. The slides were subsequently processed according to the guidelines provided in the CosMx SMI Slide Preparation for FFPE RNA Assays manual (NanoString, MAN-10184-02), starting from page 38. Slides were immersed directly in pre-heated 1x Target Retrieval Solution (NanoString, CosMx FFPE Slide Preparation RNA Kit) in a pressure cooker at 100 °C for 8 min (as per MAN-10184-02). The slides were immediately transferred to water for 15 s, then washed in 100% ethanol for 3 min, and air-dried at RT for 30 min to 1 h. Incubation frames were attached to each slide, and a pre-warmed digestion buffer containing 1.5 µg/mL Proteinase K (NanoString, CosMx FFPE Slide Preparation RNA Kit) and 1x PBS (ThermoFisher Scientific, AM9625) was applied. Slides were incubated in a hybridization chamber at 40 °C for 15 min. The slides were then rinsed twice in water, and fiducials were applied at 0.001%, followed by a 5 min incubation at RT, shielded from light. The slides underwent a 1x PBS wash for 1 min, followed by fixation in 10% NBF for 1 min, and then two washes in NBF stop buffer (Tris Base, Sigma S6639-1L, and Glycine, Sigma, G7126) for 5 min each, and a 5 min wash in 1x PBS. A 100 mM NHS-acetate solution (ThermoFisher Scientific, 26777) was applied to the tissue for 15 min at RT, followed by two washes in 2X SSC (ThermoFisher Scientific, AM9763) for 5 min each. The CosMx Human Universal Cell Characterization RNA Panel (NanoString, CMX-H-USCP-1KP-R), targeting 950 human genes, and a 50-target add-on panel set were denatured at 95 °C for 2 min, cooled on ice for 1 min, and then added to a probe mix containing RNase inhibitor, Buffer R, and nuclease-free water. This mix was applied to the slide and incubated for 18 h in the hybridization chamber at 37 °C. After incubation, the slides were washed twice in a final concentration of 50% deionized formamide (ThermoFisher Scientific, AM9342) and 2X SSC mix for 25 min each, followed by two washes in 2X SSC for 2 min each. DAPI nuclear stain stock was diluted to 1:40 with blocking buffer (Nanostring, CosMx FFPE Slide Preparation RNA Kit) and applied to the slides for 15 min at RT, protected from light. The slides were then washed in 1x PBS for 5 min and stained for 1 h with a cocktail of CD298, B2M, PanCK, and CD45 antibodies (Nanostring, 121500020, 121500021). The slides were washed three times in 1x PBS for 5 min each and then stored in 2X SSC. The pre-bleaching profile followed configuration A, while the cell segmentation profile adhered to configuration C (MAN-10161-03-2, Nanostring).

### CosMx Protein Slide Preparation (STAMP-X-CP/CP)

STAMPed slides (for both single and multimodal STAMP) were processed according to the guidelines provided in the CosMx SMI Slide Preparation for FFPE Protein manual (MAN-10185-01-1, NanoString) from page 31. Briefly, slides were baked at 65 °C for 2 h, equilibrated to RT for 3–5 min, then rehydrated in 1x PBS for 5 min. Slides were immersed in pre-heated 1x Target Retrieval Solution (Nanostring, CosMx Protein Slide Preparation FFPE Kit) in a pressure cooker at 100 °C for 8 min and then allowed to equilibrate to RT in the same solution for 60 min. Subsequently, the slides were washed three times in 1x PBS for 5 min each, and incubation frames were attached. Slides were then covered with Buffer W and incubated at RT for 1 h in the dark. The 64-plex human immuno-oncology protein antibody panel (Nanostring, CMX-H-IOP-64P-P) was combined with CD298/B2M and PanCK/CD45 segmentation markers in Buffer W. The primary antibody mix was incubated at 4 °C for 18 h, followed by three washes with 1x TBS-T buffer for 10 min each and a wash with 1x PBS for 2 min. Fiducials, prepared at the recommended concentration of 0.00005%, were applied to the slides at RT for 5 min, protected from light. Slides were then washed once with 1x PBS for 5 min and fixed with 4% PFA for 15 min, followed by three washes in 1x PBS for 5 min each. Sections were stained with a 1:40 diluted nuclei stain for 10 min, washed twice with 1x PBS for 5 min, and incubated with 100 mM NHS acetate for 15 min, before a final wash in 1x PBS for 5 min. The selected pre-bleaching profile was Configuration A, and the cell segmentation profile was Configuration C.

### CosMx SMI Setup and Data Acquisition (STAMP-X-CP/CP/C)

The setup and scan acquisition for the CosMx SMI instrument were performed according to the CosMx SMI Instrument User Manual (MAN-10161-03-2, Software Version 1.3.0.209, NanoString). A new acquisition process was initiated through the CosMx SMI Control Center web interface. Before insertion into the CosMx Flow Cell Assembly Tool, the slides were carefully dried in the areas surrounding the imaging region. The assembly process involved lowering the tailgate, placing the slide, and securing it by raising the tailgate. After removing the adhesive backing, a new flow cell coverslip was precisely aligned above the slide’s imaging area. The Assembly Tool’s lid was then closed securely to attach the coverslip to the slide, creating a functional CosMx flow cell. Following assembly, 2X SSC (for RNA assay) or 1x PBS (for protein assay) was gently introduced through one of the flow cell ports to hydrate the tissue section. Flow cell configuration data, including the flow cell’s barcode, slide ID number, and the maximum tissue thickness of 7 µm, were entered into the Control Center interface (MAN-10161-03-2, Nanostring). All slides were scanned using Configuration A for the pre-bleaching profile and Configuration C for the cell segmentation profile. Additional details regarding the probe panel, cell segmentation, and supplemental markers were also entered into the flow cell configuration data for each section. Assembled flow cells/slides, along with Buffer Bottles 1-4, were loaded into the instrument. Before positioning Bottle 4, catalase and pyranose oxidase enzymes were added. For RNA runs, an RNase inhibitor was added to a designated well in the CosMx imaging tray, which was placed in the instrument after equilibration to RT. The Control Center configuration was verified, followed by a pre-run check and a tissue find scan for each slide conducted by the instrument. After completing the tissue find scans, rectangular scan areas were placed around each tissue section for the preview scan. The preview scan images enabled the selection of regions of fields of view (FOVs) using the grid FOV placement tool, ensuring thorough coverage of the STAMP areas on each slide. FOV selections for each slide were confirmed before starting the cycling process. CosMx scan data was automatically uploaded to NanoString’s cloud-based AtoMx Spatial Analysis Platform during the run, as detailed in the CosMx Data Analysis Manual (MAN-10162-03, Software Version 1.3.2, NanoString). Upon completion of the run and full upload of scan data to AtoMx, a study was created for each CosMx scan. Within AtoMx, pipelines were executed for each study, and data was exported in various formats, including TileDB arrays and Seurat objects, for in-house analysis.

### Xenium Slide Preparation (STAMP-X)

STAMPed Xenium slides were placed on a Xenium Thermocycler Adapter plate atop the 96-well block of a thermocycler with the lid open and incubated at 60 °C for 2 h for Xenium v1 or 30 min for Xenium Prime. Slides were then equilibrated to RT for 7 min, assembled into Xenium cassettes, and hydrated with 1x PBS for Xenium v1 or PBS-T for Xenium Prime. The slides were processed following the Xenium In Situ for FFPE Deparaffinization and Decrosslinking protocol from step 1.4.a (page 42) (CG000580 Rev D, 10X Genomics). Briefly, the slides were reverse crosslinked using a decrosslinking buffer containing tissue enhancer, urea, and perm enzyme B at 80 °C for 30 min, followed by three washes with PBS-T. For Xenium v1, slides were then immediately processed according to the Xenium In Situ Gene Expression Cell Segmentation User Guide (10X Genomics, CG000749 Rev A) for the remaining slide preparation steps. Briefly, the pre-designed gene expression probe set, Xenium Human Immuno-oncology Panel (10X Genomics, PN-1000654), targeting 380 human genes, was denatured at 95 °C for 2 min, crash-cooled on ice for 1 min, and then equilibrated to RT before being added to the probe hybridization buffer and TE buffer to make the probe hybridization mix. The slides were incubated with the hybridization mix at 50 °C for 20 h. Slides were washed three times with PBS-T for 1 min each and then incubated with a post-hybridization wash buffer for 30 min at 37 °C. Slides were washed three times with PBS-T for 1 min each and incubated with the ligation mix for 2 h at 37 °C. Three 1 min PBS-T washes were followed by a 2 h incubation at 30 °C with amplification mix and enzyme (10X Genomics, PN-2000392, 2000399). Slides were then washed three times with TE buffer for 1 min each, 70% ethanol for 2 min, twice with 100% ethanol, and once with 70% ethanol for 2 min each before rehydration with PBS-T. Slides were incubated at RT for 1 h with 1x Xenium Block and Stain Buffer (10X Genomics, PN-2001083). For segmentation staining, slides were incubated for 20 h at 4 °C with the Xenium Multi-Tissue Stain Mix (10X Genomics, PN-2000991), which contains a cocktail of antibodies labeling the membranes (anti-ATP1A1/CD45/E-cadherin), antibodies labeling the cell interior (anti-alphaSMA/Vimentin), and a universal interior label against Ribosomal RNA (18S rRNA) (10X Genomics, PN-2000991). Staining was enhanced by the addition of Xenium Staining Enhancer reagent (10X Genomics, PN-2000992), followed by treatment with Xenium Autofluorescence Mix (10X Genomics, PN-2000753) to diminish unwanted autofluorescence and enhance the signal-to-noise ratio. Subsequently, Xenium Nuclei Staining Buffer (10X Genomics, PN-2000762) was used to facilitate the identification of tissues or regions of interest during the instrument’s overview scan. For Xenium Prime, slides were immediately processed following the Xenium Prime In Situ Gene Expression with Optional Cell Segmentation User Guide (CG000760 Rev A, 10X Genomics) for the remaining slide preparation steps. Briefly, following decrosslinking steps in CG000580, Xenium 5K Human or Mouse PTP Panel Priming Oligos (10X Genomics, PN-2001224 or 2001226) were denatured at 95 °C for 2 min, crash-cooled on ice for 1 min, and then equilibrated to RT before being added to Priming Hybridization Mix with TE buffer and Priming Hybridization Buffer (10X Genomics, PN-001228). A Xenium Cassette Insert was placed onto the Xenium Cassette v2, and the slides were incubated with the priming hybridization mix at 50 °C for 1.5 h, then washed twice with PBS-T and incubated with Post-Priming Wash Buffer (10X Genomics, PN-2001229) at 50 °C for 30 min. Following three 1 min PBS-T washes, slides were incubated with an RNAse mix containing 2X RNAse buffer, RNase enzyme (10X Genomics, PN-2000411, 3000953), and water at 37 °C for 20 min. Slides were then washed three times with 0.5X SSC-T and incubated with a polishing reaction mix containing Polishing Buffer, Polishing Enzyme (10X Genomics, PN-2001231, 2001230), and water at 37 °C for 1 h. The pre-designed gene expression probe set, Xenium Prime 5K Human or Mouse Pan Tissue & Pathways Panels (10X Genomics, PN-1000724, 1000725), was denatured at 95 °C for 2 min, crash-cooled on ice for 1 min, and then equilibrated to RT before being added to the probe hybridization buffer and TE buffer to make the probe hybridization mix. Following three 1 min PBS-T washes, Xenium Cassette Inserts were placed onto the Xenium slide, and the slides were incubated with the hybridization mix at 50 °C for 20 h. Slides were washed twice with PBS-T for 1 min each, then incubated with a post-hybridization wash buffer (10X Genomics, PN-2000395) for 15 min at 35 °C. Slides were washed three times with PBS-T for 1 min each and incubated with ligation mix containing ligation enzymes A, B, and ligation buffer (10X Genomics, PN-2000397, 2000398, 2001233) for 30 min at 42 °C. Following ligation, slides were washed three times with PBS-T for 1 min each and incubated with Amplification Enhancement Master Mix containing Amplification Enhancer Buffer and Amplification Enhancer (10X Genomics, PN-2001234, 2001235) at 4 °C for 2 h. Amplification Enhancer Wash Buffer (10X Genomics, PN-2001236) was added, and slides were incubated with amplification mix (10X Genomics, PN-2000392) at 30 °C for 1.5 h, followed by three 1 min washes with TE buffer. Slides were processed for segmentation staining as outlined above for Xenium v1.

### Xenium Analyzer Setup and Data Acquisition

STAMPed Xenium slides, assembled within Xenium cassettes, were imaged using the Xenium Analyzer in accordance with the guidelines specified in the Xenium Analyzer User Guide (CG000584 Rev F, 10X Genomics). The Xenium Decoding Consumables Kit (PN-1000487, 10X Genomics) was used for instrument loading. Briefly, Xenium slide ID numbers and information on pre-designed gene expression probes were input into the Analyzer, and the necessary consumables and reagents were loaded into the instrument. Reagent modules B, and C for Xenium v1, were thawed at 4 °C overnight and equilibrated to RT for 30 min before loading into the instrument, while reagent module A was stored at 4 °C until loaded. Instrument wash buffer (100% Milli-Q water), sample wash buffer A (1x PBS, 0.05% Tween-20 (ThermoFisher Scientific, 28320), sample wash buffer B (100% Milli-Q water), and probe removal buffer (50% DMSO (Sigma Aldrich, D8418) 50 mM KCl (ThermoFisher Scientific, AM9640G), 0.1% Tween-20) were prepared and loaded into the instrument along with buffer caps, a pipette tip rack, an extraction tip, and the objective wetting consumable. After loading, the samples were scanned to generate images of the fluorescently labeled nuclei in each section, which were used for Field of View (FOV) selection prior to run initiation. Each STAMP area was selected as a separate region and labeled accordingly. Upon completion of the run, the instrument was cleared of consumables, and the Xenium slides were carefully removed. Fresh PBS-T was applied to each slide/cassette, which were then covered and stored in the dark at 4 °C for up to 3 days until post-run H&E staining. The data were acquired using the Xenium Explorer software suite (v3.1.0, 10X Genomics), which provides a set of applications for analyzing and visualizing in situ gene expression data produced by the Xenium Analyzer.

From Xenium RNA to CosMx Protein (STAMP-X-CP) and to PCF (STAMP-X-PCF) Following the completion of the Xenium Prime 5k Human RNA run, slides were removed from the instrument and stored in the Xenium cassette with 50% glycerol in 1x PBS at 4 °C. After 6 days of storage, one of the slides (STAMP-X-CP) was washed three times in 1x PBS for 1 min each, followed by two additional washes in 1x PBS for 5 min each. The Xenium cassette was then removed, and the slide was placed into a wash jar containing 1x PBS for 5 min, as described in the CosMx SMI Manual Slide Preparation for Protein Assays (MAN-10185-01-1, NanoString). The slide was then immersed in antigen retrieval solution and incubated for 8 min at 100 °C, according to the CosMx Protein Slide Preparation and CosMx SMI Setup and Data Acquisition protocols. The other slide (STAMP-X-PCF) was covered with 50% glycerol in 1x PBS, coverslipped and sealed for shipment to The Jackson Laboratory (CT, USA) for PhenoClycler Fusion profiling (see below).

PhenoCycler Slide Preparation (STAMP-X-PCF/PCF) The STAMP-PCF slide was baked at 60 °C for 30 min. During the final 5 min of baking, the STAMP-X-PCF slide was removed from 50% glycerol/PBS storage and placed in 1x PBS. After baking, both slides were washed twice in 1x PBS for 5 min each. The slides were then immersed in 1x antigen retrieval buffer (pH 9.0) (AR9, Akoya Biosciences), and antigen retrieval was performed at 95 °C for 8 min using the TintoRetriever (BioSB). Following antigen retrieval, the slides were cooled in the retrieval buffer to RT and washed in nuclease-free water for 5 min. Slides were then processed according to the PhenoCycler-Fusion User Guide (PD-000011 REV M, Akoya Biosciences), starting from step 4 on page 49. Briefly, the slides were washed in Hydration Buffer, equilibrated in Staining Buffer, and incubated overnight at 4 °C with a 40-marker antibody cocktail prepared in Blocking Buffer. The slides were subsequently washed in Staining Buffer, gently fixed with Post-Staining Fixing Solution, washed in 1x PBS, and incubated in ice-cold methanol for 5 min. The slides were then washed in 1x PBS, fixed with Final Fixative Solution for 20 min, washed three times in 1x PBS, and immersed in Storage Buffer prior to the PhenoCycler Fusion run.

### PhenoCycler Setup and Data Acquisition (STAMP-X-PCF/PCF)

The experimental protocol was set up using the PhenoCycler Experiment Designer (Version 2.1.0, Akoya Biosciences). A reporter plate containing fluorescently labeled barcode reporters, as per the experimental design, was prepared following instructions on page 73 of the PhenoCycler-Fusion User Guide (PD-000011 REV M, Akoya Biosciences). Slides were prepared for the PhenoCycler-Fusion instrument according to the steps outlined in the PhenoImager-Fusion User Guide (PD-000001 Rev N, Akoya Biosciences). Briefly, the slide was moved from Storage Buffer to 1x PBS and incubated for 10 min. After incubation, a Flow Cell (Akoya Biosciences) was attached to the sample slide using the Flow Cell Assembly Device (Akoya Biosciences). The slide with the attached Flow Cell (Sample Flow Cell) was then placed in 1x PhenoCycler buffer for 10 min. PhenoCycler Fusion software (Version 2.2.0) was used to set up the imaging run on the PhenoCycler-Fusion, following the steps on page 57 of the PhenoImager-Fusion User Guide (PD-000001 Rev N, Akoya Biosciences).

Reagents were prepared and loaded into the appropriate reagent reservoirs on the instrument, and the pre-prepared reporter plate was loaded into the PhenoCycler. A new PhenoCycler run was initiated using the experimental protocol design. A blank flow cell was loaded into the Flow Cell Slide Carrier, and all software prompts during the pre-flight routine were followed. The Sample Flow Cell was then loaded into the carrier, and a leak check was performed. Scan regions were selected following automated sample finding, and imaging was started. Upon completion, the Sample Flow Cell was placed in Storage Buffer at 4 °C. The generated QPTIFF data file was used for downstream image analysis.

### H&E Staining and Imaging

For post-Xenium (STAMP-X) Hematoxylin and Eosin (H&E) staining, slides were quenched in 10 mM sodium hydrosulfite (Sigma Aldrich, 157953-5G) at RT for 10 min, rinsed three times in water, and then immediately processed through the following sequence: once in water for 2 min, once in Mayer’s Hematoxylin (Sigma Aldrich, MHS16) for 20 min, three times in water for 1 min each, once in bluing solution (Dako, CS702) for 1 min, once in water for 1 min, once in 70% ethanol for 3 min, once in 95% ethanol for 3 min, once in Eosin Y Solution, Alcoholic (Leica, 3801615) for 2 min, twice in 95% ethanol for 30 seconds each, twice in 100% ethanol for 30 seconds each, and twice in xylene for 3 min each, as described in the Demonstrated Protocol Xenium HE Staining (CG000613 Rev B, 10X Genomics). The slides were dried for 15 min and then cover slipped using 1.5 mm thick cover glass and Cytoseal Mountant XYL (ProSciTech, 1A013-XYL-118) or toluene-free mounting media (Dako, CS705). For post-CosMx (STAMP-X-CP/CP/C) H&E staining, the glass flow cell coverslip was first removed by adhering clear sticky tape to the top of the flow cell, scoring around the inside of the adhesive edges of the flow cell, then peeling off the sticky tape, which left the adhesive edges attached to the slide and exposed the tissue for staining. Slides were then washed by dipping into water several times to remove any glass shards. The Demonstrated Protocol Xenium HE Staining (CG000613 RevB, 10X Genomics) was then followed starting from step 1.4, as described for post-Xenium H&E staining. Stained sections were covered using custom-cut coverslips fitted inside the flow cell adhesive edges using a glass scribe. Once the mounting media had dried, slides were scanned using a NanoZoomer 2.0HT (Hamamatsu) with a 40X objective or a Zeiss AxioObserver 7 with a 20x objective (Zeiss). For post-PhenoCycler (STAMP-X-PCF/PCF) H&E staining, the Sample Flow Cells were removed from the Storage Buffer and placed in a coplin jar containing xylene for 24 h. The Flow Cells were then carefully removed and disposed of properly. The slides were transferred to 100% ethanol for 2 min, dipping 10-15 times to ensure full coverage of the tissue. This step was repeated using 95% ethanol, followed by DI water. The slides were then placed in Mayer’s Hematoxylin for 4 min, followed by a rinse in DI water for 1 min, Bluing Reagent for 1 min, another DI water rinse for 1 min, and stained in Alcoholic Eosin for 2 min. Subsequent washes included 95% ethanol for 1 min, 100% ethanol for 1 min, fresh 100% ethanol again for 1 min, xylene I for 1 min, and finally, the slides were held in fresh xylene. Inside a fume hood, each slide was removed one at a time, the back was dried, and the slide was tilted onto a paper towel to remove excess xylene without allowing the tissues to dry. DPX or another xylene-based mountant was applied over the tissue area using a disposable Pasteur pipette while the remaining slides were kept in xylene to prevent over-drying. The glass coverslip was swiftly cleaned, any particles were removed, and the long edge was placed onto the slide. The slide was tipped towards the user, allowing the mountant to contact the coverslip, and was gently pressed down until the mountant spread evenly across the tissue. Excess mountant was carefully blotted using a paper towel, avoiding contact with the top of the coverslip. Any bubbles over the stained tissue were gently pressed out. The mountant was allowed to cure in the hood for at least 20 min before imaging. The slides with coverslips were then imaged using the NanoZoomer-SQ Digital slide scanner (Hamamatsu) with a 40x objective.

### Chromium Fixed RNA Profiling of Cancer Cell Lines and hESC and Sequencing

MCF-7, LnCAP, SK-BR-3, iPSC, and hESC cell lines were processed using the Fixation of Cells & Nuclei for Chromium Fixed RNA Profiling protocol (CG000478, RevD, 10X Genomics). After quenching, the cells were counted in replicates using the Luna-FX7 Automated Cell Counter, and 0.5 to 1 million cells were subjected to either the Chromium Fixed RNA Kit (PN-1000474, 10X Genomics) for cancer cell lines or the Chromium Fixed RNA Kit (PN-1000475, 10X Genomics) for iPSC lineages and hESC BMP4 treatments. Singleplex or multiplex gene expression libraries were prepared according to the user guides CG000691 (RevB) and CG000527 (RevF), respectively. The libraries were quality controlled using a 5200 Fragment Analyzer System (Agilent, HS NGS Fragment Kit, DNF-474-1000) and sequenced on a NovaSeq 6000/X instrument following 10X Genomics’ user guide recommendations.

## Data Analysis

### Fixed RNA Profiling (Flex)

The Flex datasets were aligned to probe set reference from 10X Genomics using cellRanger v8.0 with the filter-probes argument set to false. Filtered feature-barcode matrices were loaded into *R* (v4.4.1) as a *SingleCellExperiment* object using the *read10xCounts* function from the *DropletUtils* package (v. 1.24.0)^20,21^.

### Quality Control

Quality control metrics were computed for each cell using the *addPerCellQC* function from the *scater*^22^ package (v1.32.0). Outliers in the number of counts and detected features were identified and removed using the *isOutlier* function from *scater* with parameters *type = “lower”*, *log = TRUE,* and *nmads = 2* for counts and *nmads = 2.5* for detected features, respectively. Cells with a high percentage of mitochondrial gene expression were filtered out by applying *isOutlier* with *type = “higher”*, l*og = FALSE*, and *nmads = 5*.

### Pre-processing

The count data were log-normalized using the *logNormCounts* function from *scater*. Feature selection was performed by modeling the mean-variance relationship with the *modelGeneVar* function from the *scran*^23^ package (v1.32.0). Highly variable genes were selected using *getTopHVGs* with an FDR threshold of 0.7. Principal Component Analysis (PCA) was conducted using the *fixedPCA* function from *scran*, specifying the *subset.row* argument to include the selected highly variable genes. The first 25 PCs were selected for downstream analyses based on the variance explained. UMAP dimensionality reduction was then applied using the *runUMAP* function from *scater* on these components.

### Clustering

Cells were clustered using the *clusterCells* function from *scran*, specifying *dimred = “PCA”* and *BLUSPARAM = NNGraphParam(k = 50, cluster.fun = ”louvain”)*.

### Doublet Identification

Potential doublets were identified using the *scDblFinder* function from the *scDblFinder* package^24^ (v1.16.0), incorporating cluster information via the *clusters = colLabels(sce)* argument, and subsequently removed by looking at the *scDblFinder.score* metrics together with canonical markers exclusive of specific populations.

## CosMx RNA Datasets

### Cell segmentation in AtoMx

Default cell segmentation yielded suboptimal results (e.g. over-segmentation and / or over-expanded cell boundaries) as assessed by visual inspection of the immunofluorescence images (e.g. DAPI, panCK, CD45 and CD298/B2M). Thus, we re-run cell segmentation for all STAMP-C experiments within the AtoMx platform using *Configuration C* (*Cell Pellet Array*) and setting the *Basic Parameters* as follows: *CellDilation* at 2 µm, *CellDiameter* at 30 µm for cell lines and 10 µm for PBMCs and *NuclearDiameter* at 10.8 µm for cell lines and at 5 µm for PBMCs. We additionally adjusted the *Advanced Parameters* for STAMP-C shown in Fig. 1 and Suppl. Fig. 1 as follows: *BackgroundPercentile* at 0.4, *LogBlurSigma* at 2, *LoGThreshold* at 5, *Nuclei* and *CytoplasmModels* at CP, *NucleiProbability* at -2, *CellProbability* at -3, *CellFlowThreshold* at 0.1 and *MinCellSize* at 4.3. Flat-files exported from AtoMx were imported into R (v4.4.1), generating SingleCellExperiment objects using the SingleCellExperimens package^25^ (v1.26.0).

### Quality Metrics

For each cell, we considered the cell area, number of counts and number of detected genes (the latter computed using the *addPerCellQC* function from the *scater* package (v1.32.0)). Lower and Higher outlier cells in any of these distributions were identified and removed using the *isOutlier* function from *scater* with the parameter *log=TRUE*. The *nmads* parameter, which specifies the number of median absolute deviations for outlier detection, was determined independently for each dataset based on visual inspection of the distributions. Additionally, cells located within 30 pixels of any field of view border were filtered out.

### Clustering

Cells were clustered using the unsupervised approach provided by the *InSituType* (IST)^16^ package (v1.0.0), with the *n_clust* parameter varying according to the annotation step.

Background noise was calculated using the mean negative probe counts. The *fastCohorting* function was used to perform cohorting on mean immune fluorescence values (CD298/B2M, CD45, PanCK, DAPI, CD68_CK8_18), cell area, and aspect ratio. Cells with a posterior assignment probability below 0.8 were excluded from further analysis.

### Marker genes

IST generates cluster profiles by aggregating counts for each gene within each cluster, and correcting for background (mean negative probe count). To compute marker genes, we normalized these cluster-level counts using the *normalizeCounts()* function from the *scater* package, and calculated the log2 fold change between each cluster and the average across remaining clusters (adding a small constant of 1e-6 to both).

### UMAP

Prior to dimensionality reduction, the count matrices were normalized by total counts and log1p-transformed. Principal Component Analysis (PCA) was performed using the *prcomp_irlba* function from the *irlba* package^26^ (v2.3.5.1), a subset of PCs chosen based on the elbow plot, and Uniform Manifold Approximation and Projection (UMAP) was applied using the *umap* function from the *uwot* package^27^ (v0.2.2).

### Signature Scoring

The circulating tumor cell mimics signature was derived by subsetting a 10x Genomics Flex dataset containing peripheral blood mononuclear cells (PBMCs) and MCF-7 cells, retaining only genes present in the CosMx 1K panel. We then applied the *scoreMarkers* function from the *scran* package (v1.32.0) to identify marker genes. The top 100 markers were then selected by highest absolute log2FC and the signature was subsequently scored in the PBMCs-CTCs CosMx dataset using the *AUCell_run* function from the *AUCell* package^28^ (v1.26.0).

### Differentiation trajectory

After the preprocessing of the hESC samples, we used more strict thresholds to remove low quality cells. More specifically, we removed cells with more and fewer than 300 and 600 genes and 1500 and 6000 counts, respectively. Then, for each time point, we normalized and scaled the data again using the NormalizeData() and ScaleData() functions of Seurat (version X) followed by a principal component analysis and clustering using 30 principal components and a clustering resolution of 0.1. As we expected the distinct cell states to be uniformly distributed across the FOVs, we decided to remove cell clusters that were present in less than half of the FOVs. To create a differentiation trajectory we used the Palantir algorithm on a subset of ∼40,000 cells. Briefly, Palantir constructs an augmented affinity matrix by augmenting the kNN graph affinity matrix with mutually nearest neighbors between successive time points. This matrix forms the basis to generate a force directed layout for visualization and as input for computing the diffusion operator which can be used for trajectory detection. To generate the Amnion, Endoderm and Mesoderm trajectories we determined the cell of origin and the terminal cells for each trajectory.

## CosMx Protein Datasets

Cell segmentation and initial pre-processing were performed using AtoMx. Flat files were downloaded, and background fluorescence was subtracted from each segmented cell to control for non-specific fluorescence signals. Cells with aggregated signals less than 20 or greater than 10,000, as well as those with an area smaller than 40 um², were excluded from further analysis. Seurat was used for dimensionality reduction and clustering, selecting all 42 proteins for analysis. Data was normalized by dividing the fluorescence intensity for each cell by the total intensity for that cell, multiplying by 1e6, and then natural-log transformed. Principal component analysis was performed with nPCs set to 40, and after examining the elbow plot, the top 15 PCs were chosen for constructing the neighborhood graph (sNN, k.param = 20, distance = “euclidean”). Louvain clustering was then conducted with a resolution of 0.8. Cell annotation was achieved by inspecting the expression of each protein in the panel across the identified clusters, resulting in the classification of cells into eight main categories. For quality control of cells profiled in CosMx Protein after Xenium and those profiled in CosMx Protein alone, aggregated expression was calculated for each protein. Using the R packages ggpubr and ggplot2, a scatter plot was generated to compare the average expression of each protein across the different experiments. A linear regression line was added to represent the best-fitting line through the data points, along with Pearson correlation statistics to assess the relationship between the datasets.

## Xenium Datasets

### Quality Metrics

Low-quality cells were filtered out by applying data-driven adaptive thresholds to the distributions of counts, genes, and cell area, following the same approach as with the CosMx datasets.

### Pre-processing

SingleCellExperiment objects of the Xenium datasets were log-normalized using the l*ogNormCounts* function from the *scuttle* package (v1.14.0). For the Immune-Oncology datasets, no feature selection was performed. PCA was conducted using the *fixedPCA* function from *scran* with the parameter *BSPARAM=IrlbaParam()*. The proportion of variance explained by each principal component guided the selection of components for downstream analyses. UMAPs were then generated using the *runUMAP* function from the *scater* package.

### Clustering

We constructed a shared nearest neighbors (SNN) graph using the *buildSNNGraph* function from *scran*, specifying *type=“jaccard”,* k = 50, and *BNPARAM=AnnoyParam()*. Louvain clustering was performed on this graph using the *cluster_louvain* function from the *igraph* package^29,30^ with resolutions 0.5 or 1 depending on the annotation step. Following the initial round of annotation to identify major lineages, a second round of pre-processing and clustering was conducted within these lineages to achieve more detailed annotation.

### Feature Selection

Following an initial annotation round—conducted by applying the aforementioned steps to the entire panel— and only in the 5K panel datasets, we performed feature selection to identify highly variable genes (HVGs) within PBMC lineage subsets. The mean-variance relationship was modeled using the *modelGeneVar* function from *scran*, and HVGs were selected using the *getTopHVGs* function setting the *fdr.threshold = 0.9*. PCA was then rerun as previously described, incorporating the *subset.row* parameter to specify the selected HVGs.

### Akoya PhenoCycler Datasets

Cell segmentation was performed using StarDist in QuPath (v0.5.0) with a custom Groovy script. The following parameters were used for the StarDist function: threshold = 0.5, channel = 0, normalizePercentiles = (1, 99), pixelSize = 0.5, cellExpansion = 5.0, and cellConstrainScale = 1.5. Fluorescence measurements were exported in text format. An expression matrix was constructed, retaining only the mean intensity measurements per channel for each segmented cell. Seurat (v5.1) (10.1038/s41587-023-01767-y) was used for quality control and downstream analysis. Cells with a mean aggregated intensity below 200 or an area smaller than 20 um² were excluded. Data normalization was performed using the centered-log-ratio option of the NormalizeData function in Seurat. All 40 proteins were included in the calculation of 40 principal components, and after reviewing the elbow plot, the top 15 PCs were selected for calculating the neighborhood graph. Louvain clustering was then applied with a resolution of 0.5 for unsupervised clustering.

## Code Availability

The code used in this project can be accessed at https://github.com/emanuelepitino/Stamp_V4

## Authors contributions

L.G.M. and J.P. devised the STAMP approach, workflow, and protocols. L.G.M., H.H., A.P.R., and J.P. designed the study, coordinated the execution of all experiments and computational analyses, interpreted the data, and wrote the manuscript. E.P. led the analyses in this study and, together with F.S.D., H.L.C and I.S.M. conducted the computational work. E.P., A.P.R, F.S.D and J.N. interpreted the data. K.W., H.C., S.R., G.C. and L.G.M. performed all STAMP-X/X-CP, STAMP-C/CP, and RNA Flex experiments. A.P.R., E.P., F.S.D. an I.S.M. generated all the figures. S.S. and E.S. conducted the STAMP-X-PCF/PCF experiments and authored the relevant Methods section. E.C. and B.F. coordinated the STAMP-X-PCF/PCF experiments. A.S., K.D. and G.A.M.R. conducted the differentiation experiments. M.M., Y.L. and H.L.C. contributed to and provided guidance on the computational analyses and code development. M.M. also coordinated the metadata and repository upload. J.M.P. and M.M. critically reviewed the workflow, experimental design, and manuscript. K.W. and M.M. designed and generated the STAMP workflow figure.

## Acknowledgements

We extend our gratitude to the South Australian Genomics Centre for facilitating our access to Xenium and managing our bookings. We also thank Milenium Sciences for their unwavering support in ensuring our reagents are always delivered on time. Special thanks to Claire Penney and Brad Duncan from 10X Genomics for providing continuous technical and sales support to our team. We are grateful to Thomas Churchward, Senior Field Service Engineer at Bio-Strategy, for his exceptional 24/7 support during our runs, and to Chin Ching Chang from the NanoString support team for assistance during troubleshooting.

## Supplementary Information

**Supplementary Figure 1.**
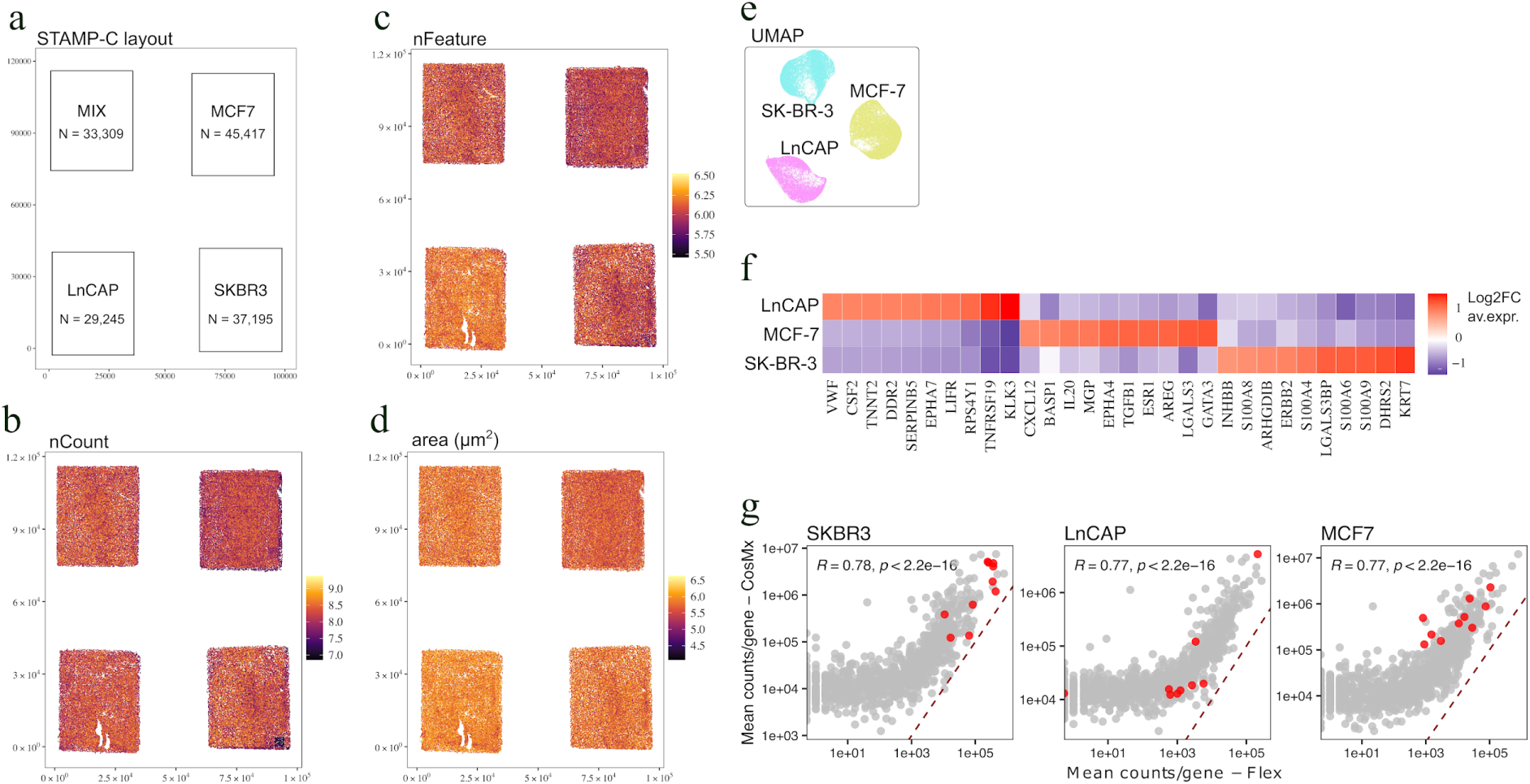
Clustering analysis of tumor cell lines in STAMP-C recapitulates classical single cell transcriptomics data. (a) STAMP-C experimental design. Spatial distribution of counts (b), features (c) and cell area (d) for each sub-STAMP. (e) UMAP color coded by cluster ID of MCF-7, LnCAP and SK-BR-3 sub-STAMPs pooled together. (f) Heat map showing the transcriptional profiles of each cell line as defined by InSituType (IST). (g) Spearman Gene expression correlation of the 10X Flex dataset and the STAMP-C from the same cell suspensions. Each dot is a gene and red dots represent marker genes for each cell line as in (f).

**Supplementary Figure 2.**
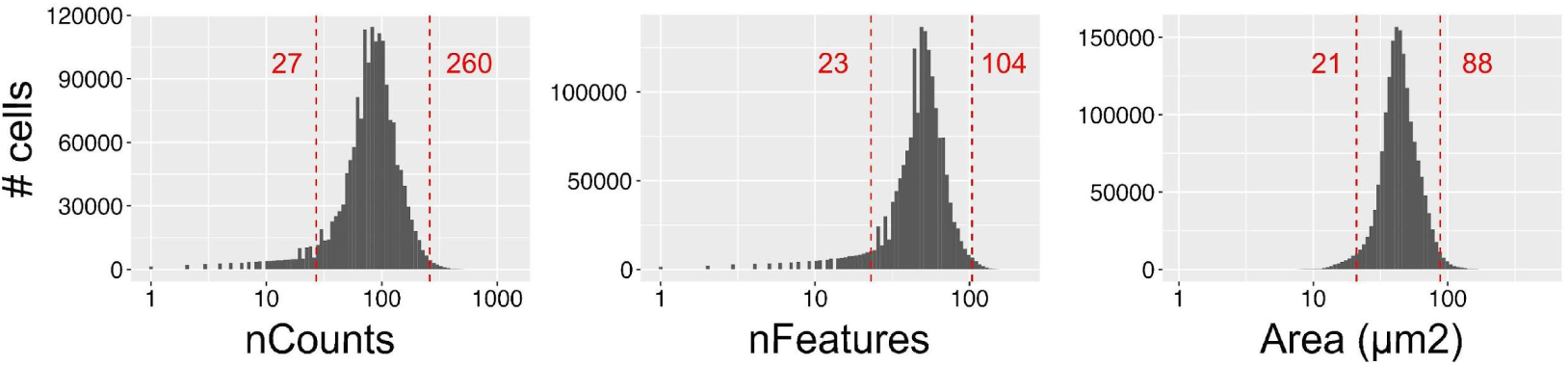
Quality control of STAMP-X data. Quality metrics of the 1.7M PBMC STAMP-x dataset showing the distribution of counts, features, and cell area before filtering. Red dotted lines and red text indicate the threshold set for filtering.

**Supplementary Figure 3.**
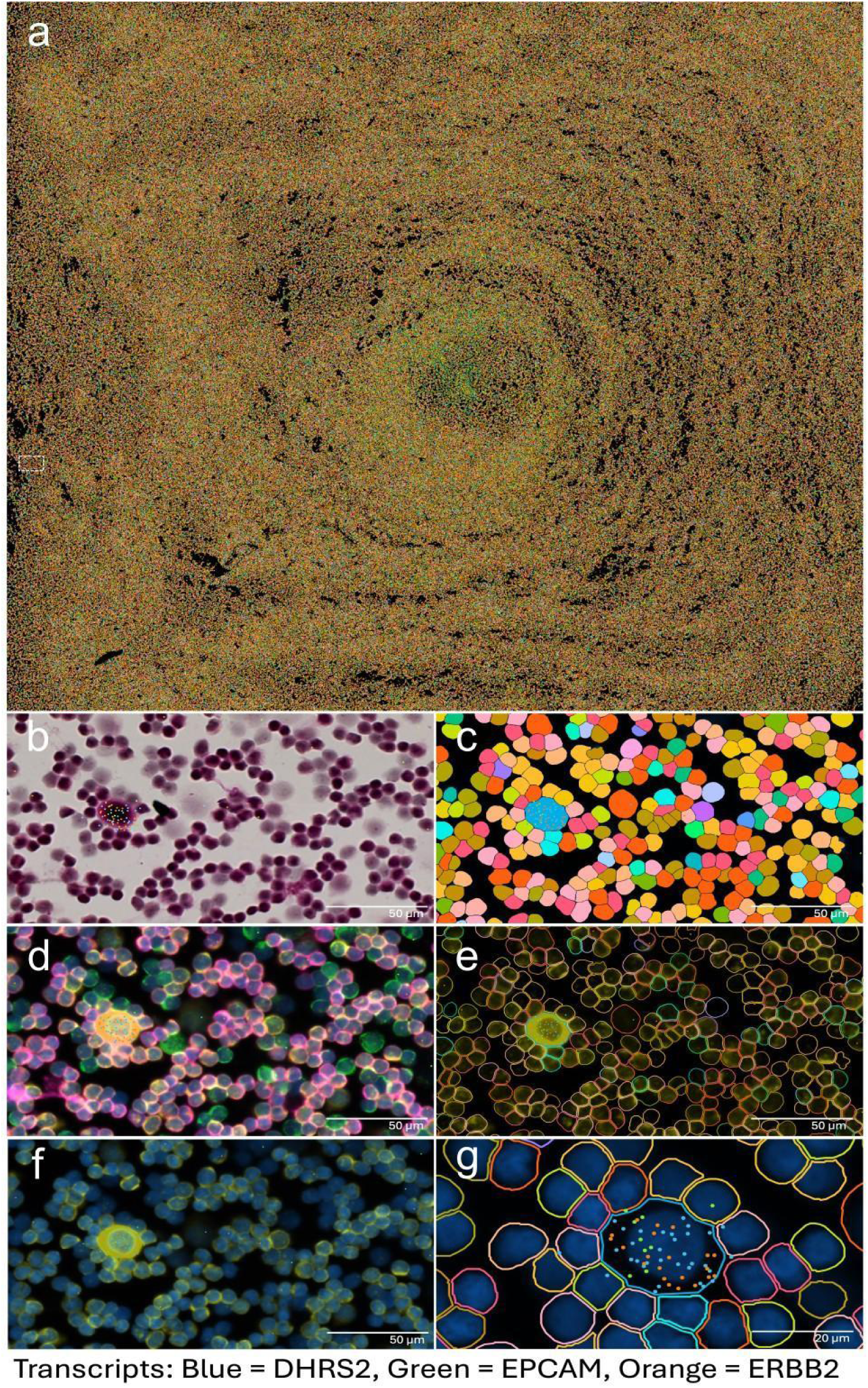
Identification of 1 CTC-mimic with STAMP-X. (a) Full stamp showing cells clustered in Xenium explorer. Region of interest is shown as white frame, from where the below images are taken from. (b) Hematoxilin and Eosin staining with single MCF-7-related transcripts. (c) Segmented cells colored by cluster ID in Xenium explorer. (d) Xenium segmentation markers: DAPI in blue, ATP1A1/CD45/e-Cadherin in pink, interior RNA (18S) in yellow, interior protein (alphaSMA/Vimentin) in green. (e) Interior RNA stain in yellow with segmentation boundaries. (f) Interior RNA stain in yellow with DAPI in blue. (g) DAPI in blue with cell boundaries colored by cluster ID and single MCF-7-related transcripts.

